# Maximum unified fatty acid signature analysis: A new approach to QFASA

**DOI:** 10.64898/2026.02.04.703822

**Authors:** Holly Steeves, Connie Stewart, Shelley Lang, Chris Field, Aaron MacNeil, Jennifer McNichol

## Abstract

Accurately estimating predator diets is crucial for understanding and predicting ecosystem changes. The desire for non-lethal but accurate diet estimation techniques has led to new approaches such as quantitative fatty acid signature analysis (QFASA), a now widely accepted method for estimating the diets of marine predators. We propose a novel alternative to QFASA, namely maximum unified fatty acid signature analysis (MUFASA), to estimate dietary proportions using fatty acid signatures. MUFASA is based on maximum likelihood estimation principles and consequently offers several theoretical advantages over QFASA, including estimates that possess desirable properties when model assumptions hold. In addition, the availability of a likelihood function enables the use of a broad range of existing methodologies that may have the capacity to address current well-known challenges associated with diet estimation via fatty acids. MUFASA and QFASA are compared using simulations based on wide-ranging diets, as well as real-life data from a captive study of harbour seals, for which diets are known. Estimates derived from QFASA and MUFASA are similar, suggesting that diet estimation in this context can potentially be viewed through an MLE framework. While not the primary focus of this work, bootstrap confidence intervals are also developed and preliminary results yield high coverage probabilities when the diet proportions are not near 0 or 1.

## 1 Introduction

For marine ecologists studying trophic structure, accurate diet compositions of predators are often of interest. Specifically, the desire for non-lethal sampling approaches has created a need for alternative diet estimation techniques (Beckmann et al., 2013). One such alternative is to use fatty acids (FAs), which are fundamental components of lipids, as trophic markers (Iverson, 1993). Unlike other dietary nutrients which are completely broken-down during digestion (e.g. proteins and carbohydrates), FAs are released from ingested lipids but are minimally degraded during digestion and are subsequently taken up into predator tissues in a predictable way with little modification (Iverson et al., 2004). As a result of biochemical limitations, only a relatively limited number of FAs can be synthesized by animals, therefore, it is possible to distinguish between FAs of dietary and non-dietary origins within animal tissues. As such, FAs have been used to quantitatively estimate the diet composition of predators by comparing the FA signature (vector of proportions of FAs) of the predator to the FA signatures of various potential prey while taking into account modifications due to lipid metabolism in the predator (Quantitative Fatty Acid Signature Analysis, QFASA; Iverson et al. (2004)).

QFASA uses a distance minimization algorithm between the FA signature of the predator and a linear combination of mean FA signatures of the potential prey. Because of the compositional nature of both FA signatures and the diet vector (the elements represent proportions of a whole), standard multivariate statistical analyses, including Euclidean distance, are unsuitable (Aitchison, 1986). Originally, transformations based on log-ratios were proposed to bring the compositions into real space, but FA data often include zeros, making both ratios and logarithms impractical. QFASA is able to accommodate the compositions by including a choice of appropriate distance measures. Aitchison’s distance is the recommended approach to measure distance between the compositional vectors in QFASA as it yields diet estimates with the least bias, and smallest root mean squared error (Bromaghin et al., 2015). The Kulback-Leibler (KL) distance is the most commonly used measure of distance in QFASA in practice, despite lacking properties such as scale invariance and subcompositional dominance (Johnson and Samuel, 1983). While both the Aitchison and KL measures of distance require imputing the zeros prior to their use, the chi-squared distance (Stewart, 2017) can handle zeros directly.

Although QFASA is widely used and accepted, it has some drawbacks. One issue not addressed by QFASA is that prey that are consumed by the predator are not the same individual prey sampled for the prey database. Therefore, the sample of prey may not be an accurate representation of the consumed prey. Additionally, QFASA omits variability in the prey FA signatures, as traditionally the model uses only the mean prey FA signature (see Meynier et al. (2010) for an application that uses individual prey signatures). Summarizing the prey signatures using the mean could be problematic in some cases if the mean signature does not resemble any FA signature of the species due to the presence of diverse groups within the species. Lastly, QFASA does not require any parametric distributional assumptions and therefore the algorithm inherits the commonly recognized drawbacks of nonparametric methods such as lower statistical power, limited or challenging applicability to complex designs and the preclusion of likelihood-based methodology.

In this paper, we lay the foundation for a novel alternative approach to estimating the diet of predators using FAs, based on the technique of maximum likelihood (ML) estimation and termed maximum unified fatty signature analysis (MUFASA). In MUFASA, the true consumed prey FA signatures are modelled as unobserved random effects and the sampled prey provide information about the distribution of these unknown FA signatures. The likelihood is formed by assuming that the FA signatures, transformed by the isometric log-ratio (ilr) transformation (the recommended transformation for compositional data; see Egozcue et al. (2003)) follow a MVN distribution, allowing prey FA signatures to vary about the mean. While not the primary focus of this work, a straightforward bootstrap confidence interval (CI) algorithm is also developed that computes individual confidence intervals for the true diet of each predator in a sample.

While a downside of MUFASA is that it is more complex than QFASA, our results suggest that the method produces comparable estimates, but with several advantages. First, provided model assumptions hold, estimates derived from MUFASA possess properties of ML estimators, including, but not limited to, consistency and asymptotic normality. Furthermore, MUFASA addresses many of the aforementioned practical limitations of QFASA through parametric modelling of the FA signatures and the use of random effects to account for predators not eating from the sampled prey database. Perhaps most notably, MUFASA provides an important framework that has the potential to address current challenges associated with diet estimation via FA analysis through extensions of existing likelihood-based methodology. For example, robust methodology to handle deviations from model assumptions or variable (prey species) selection techniques founded on ML estimation may now be applicable to our problem at hand. In particular, the MUFASA methodology has been leveraged in McNichol (2022) to estimate calibration coefficients alongside diet estimates, in Rideout (2025) where the Akaike information criterion is used as the basis for prey selection, and in Steeves (2020) to include the addition of covariates into the estimation process.

## 2 Methods

### 2.1 QFASA

QFASA uses a distance minimization algorithm to estimate the diet proportions of predators based on the fatty acid (FA) signatures of prey species. This method also accounts for changes caused by lipid metabolism in the predator that results in the proportions of fatty acids in the predator not matching exactly those in the prey it consumed. Assume that we have FA signatures from sampled predators and a prey database that contains FA signatures from prey species believed to be part of the predators’ diet. The first step is to then adjust or “calibrate” the observed FA signatures of the predators using calibration coefficients *c*_*k*_ (described in Iverson et al., 2004). The *k*^*th*^ fatty acid in the FA signature of a predator, denoted by *Y*_*k*_, is adjusted by the CCs with the following formula:

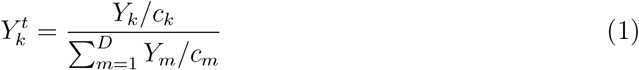

where 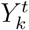 represents the CC adjusted FA, *D* is the total number of FAs included in the analysis (typically between 30 and 60) and the denominator serves to ensure the proportions sum to 1.

Let ***α*** be a vector of proportions that sums to 1, where the *i*^*th*^ element represents the proportion of prey species *i* in the predator’s diet, 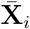 be the compositional representative FA signature of prey species *i* (usually the empirical sample mean) on the original scale, and **Y**^*t*^ be the CC-adjusted FA signature of the predator on the original scale. Then, we assume that the adjusted FA signature of the predator is approximately a linear combination of the prey means as follows:

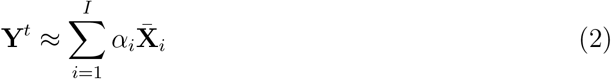

where *I* is the number of species of prey considered. In order to estimate the diet proportions ***α***, the distance between the adjusted FA signature **Y**^*t*^ and the linear combination 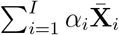. is minimized. After optimization, the lipid content of the prey is taken into account as prey species with higher lipid contents will contribute more to the FA signature of the predator than prey with lower lipid contents. The diet estimates obtained are adjusted by dividing each predator’s diet estimate by the average lipid content (% by weight) of the prey, then rescaling the diet vector to sum to 1. In the analyses presented here, calibration coefficients and lipid contents were not considered in our simulation studies (that is, they were set to vectors of 1s), but were incorporated in the real-life examples.

QFASA is commonly carried out with Aitchison or Kullback-Leibler (KL) distance. Bromaghin et al. (2015) found that Aitchison and KL distance measures yield similar results, although Aitchison’s distance yielded estimates with slightly less bias and similar or better root mean squared error (RMSE). However, in the original QFASA paper, Iverson et al. (2004) used the KL distance and this distance measure has been used in variety of validation studies (Nordstrom et al., 2008; Wang et al., 2010). Therefore, in our analysis, we will use KL distance to obtain QFASA estimates.

For readers unfamiliar with QFASA, we suggest Stewart et al. (2025), which provides an overview of FA based diet estimation techniques through examples using sample data.

### 2.2 MUFASA Diet Estimation

#### 2.2.1 Point Estimation

Building on the work of QFASA, our model involves a linear combination of diet proportions and prey FA signatures performed on the original scale, with several modifications. For simplicity, we are considering the likelihood of the *j*^*th*^ predator with diet ***α***_*j*_, and we will ignore calibration and lipid content for now. Although the FA signature of the predator is directly related to the FA signatures of the specific prey that it ate, these exact prey are unobserved. These prey FA signatures are therefore considered as random effects and integrated out of the likelihood to obtain the appropriate marginal likelihood. We can think of the predator’s FA signature as being a linear combination of the prey random effects, perturbed by some error term ***ϵ***_*j*_. Perturbation was used here since it is the simplex equivalent to addition in Euclidean space, where the simplex is the space containing all non-negative vectors that sum to 1, or the sample space of compositional data. The perturbation of a compositional vector **Z** by *ϵ* is given by **Z** *◦* ***ϵ*** = 𝒞 (*Z*_1_*ϵ*_1_, …, *Z*_*D*_*ϵ*_*D*_) where 𝒞 normalizes the composition to sum to 1. If *I* is the number of species of prey considered, then we assume

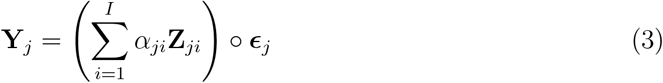

where **Y**_*j*_ is the FA signature of the *j*^*th*^ predator, *α*_*ji*_ is the proportion of the *i*^*th*^ prey species in the diet of predator *j*, **Z**_*ji*_ is the *ji*^*th*^ random effect representing the unobserved mean FA signature of prey species *i* for predator *j*, and ***ϵ***_*j*_ is the random error for predator *j* included to account for individual variability.

Since FA signatures are compositional, we utilize the isometric log-ratio (ilr) transformation (Egozcue et al., 2003) to transform the FA signatures to Euclidean space. Specifically, we used the function *ilr* in the *compositions* R package (van den Boogaart et al., 2024). A common assumption is that the ilr transformed data follow a multivariate normal (MVN) distribution in Euclidean space, hence we assume that the transformed prey (observed and unobserved) FA signatures are MVN. If the data appear more heavily skewed, then the multivariate skew normal distribution could be substituted (Azzalini and Valle (1996), Mateu-Figueras et al. (2005)). We make several further assumptions about the FA signatures of both predator and prey. We also assume that the predators’ FA signatures are conditionally independent given the random effects, since they will be sampling their own prey species. Similarly, we assume both the observed and unobserved prey are independent of each other. Since the predator is consuming the unobserved prey and has not consumed the observed prey, we can assume independence between the predator and observed prey. The unobserved prey FA signatures are assumed to be independent from each other. The observed prey FA signatures are also assumed independent from each other. Finally, we can assume that the random errors are independent of each other and are MVN after ilr transformation. Therefore, we have:

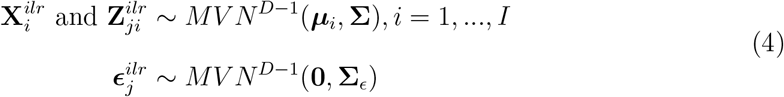

where *D* is the number of FAs, 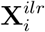 is the ilr transformation of **X**_*i*_, a randomly sampled fatty acid signature from prey species *i*, and 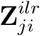 is the ilr transformed **Z**_*ji*_, the random effect representing the prey from species *i* that was consumed by the predator.

We can find the joint likelihood of 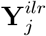 and 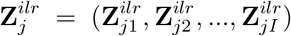 by considering the density of 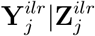 and multiplying by the density of 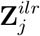. We therefore require the distribution of 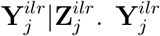 depends on 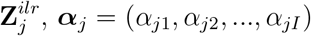 and 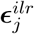. Since we are conditioning on 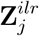, these are fixed, as are ***α***_*j*_. Therefore, the only randomness involved in 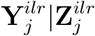 comes from 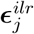. We have,

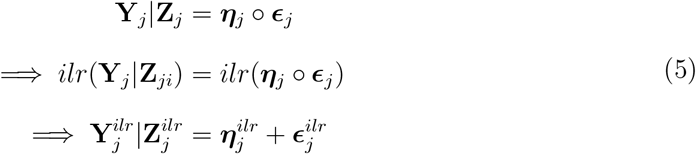

Where 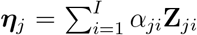. Since 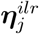 depends only on fixed ***α***_*j*_ and fixed **Z**_*ji*_, it is a constant, and 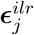 is MVN. Therefore, 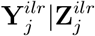 is MVN, with mean 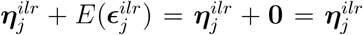. Since 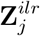 are given, the variance-covariance matrix only depends on the variance covariance matrix of the random error, **Σ**_*ϵ*_, so the variance-covariance matrix of 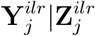 is **Σ**_*ϵ*_. Thus,

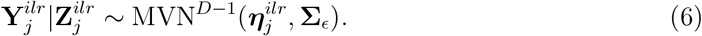

Since the random effects are unobserved, we optimize a marginal log likelihood that does not contain the 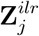. We therefore first derive the joint log likelihood and later integrate out the random effects. If we let **X**^*ilr*^ denote the matrix of all prey FA signatures and 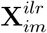 the *m*^*th*^ prey FA signature from species *i, m* = 1, …, *n*_*i*_, then from the assumptions above, we can write out the density function for the predator FA signatures, conditional on the random effects, diet proportions, variance-covariance matrix of the prey, variance-covariance matrix of the random error, the mean transformed FA signature of the prey, and observed prey FA signatures, 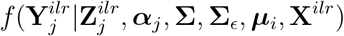. The joint likelihood can be written as

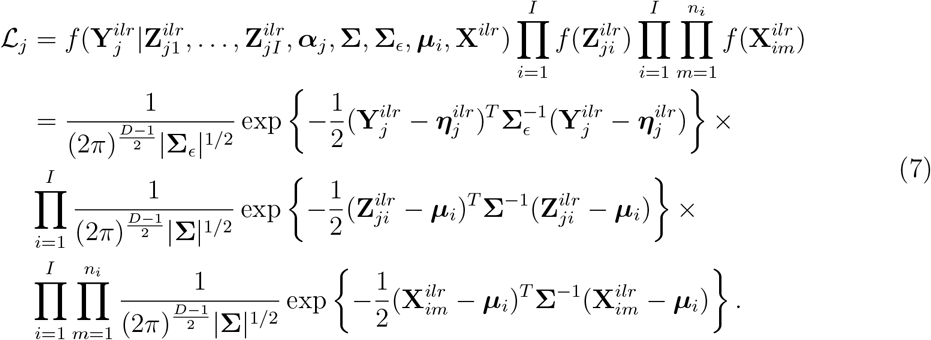

To minimize the number of parameters in the optimization, the mean ilr transformed prey FA signature, ***µ***_*i*_, is estimated using the empirical mean of the observed transformed FA signatures of prey species *i*, say 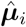, and **Σ** is estimated using the pooled empirical variance-covariance matrices of the ilr transformed prey FA signatures from the prey database, 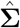. That is, a weighted average of the within prey species sample variance-covariance matrices (of the ilr transformed data) is used as the pooled estimate. We assume that **Σ**_*ϵ*_ is a diagonal matrix, with *D* − 1 values on the diagonal; however estimating all *D* − 1 values has proven difficult. Therefore, in this work, the diagonal values are grouped based on the quartiles (lower 25%, lower 25-50%, lower 50-75% and upper 25%). That is, based on the data, the diagonal elements corresponding to FAs belonging to the lower 25% are averaged, and similarly for the other quartiles. The averages are used as starting values in the algorithm and therefore only 4 distinct diagonal elements are estimated and not *D* − 1. Rather than assuming that all diagonal elements are equal or that they are all distinct, the idea of quartiles represents the reasonable assumption that the variance of the smaller FAs may be smaller than that of the larger FAs and appears to be an appropriate level of aggregation.

Notice that the last row of the likelihood in Equation 7 is the density of the prey FA signatures. When evaluated at the empirical estimates, 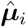, the density does not depend on ***α***_*j*_ or **Σ**_*ϵ*_, nor does it depend on the random effects 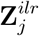. Therefore, it is a constant relative to our parameters and is not needed in the likelihood. So, the joint likelihood is

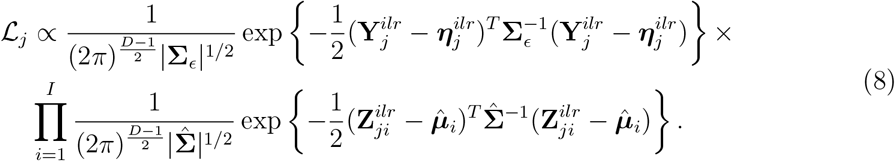

Note that the likelihood above (Equation 8) is for the *j*^*th*^ individual predator. Since the predator FAs are assumed to be independent, we can derive the joint likelihood of multiple predators simply by multiplying the likelihoods as such:

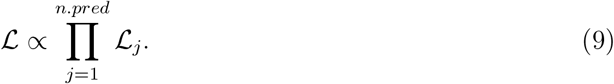

To obtain the marginal likelihood required for optimization, we need to integrate the joint likelihood in Equation 9 with respect to the random effects, 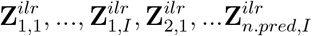. We do this using the R package *TMB* (Template Model Builder, Kristensen et al. (2016)) which approximates the marginal likelihood via the Laplace approximation. We then optimize the approximate marginal likelihood (or more accurately, the negative log of this marginal likelihood) to obtain the estimated diet proportions, 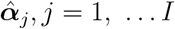. MUFASA is available as a function in the *QFASA* R package (Stewart et al., 2024).

#### 2.2.2 Interval Estimation

An attractive and well-known property of ML estimation in general is that, under certain regularity conditions and in the asymptotic setting, MLEs are normally distributed and the covariance matrix can be determined by the Fisher information matrix. The *TMB* package provides standard error estimates (see Kristensen et al. (2016) for details) and confidence intervals (CIs) based on the normal distribution follow directly for the asymptotic case.

For our application of interest, the number of parameters to be estimated tends to be large, particularly if the prey database is comprised of many different prey species. In practice, as the overall sample size may be limited and, at the predator level, the number of FAs relative to the number of prey species may be small, we would not expect estimates of the standard errors of the estimated dietary proportions or the normality assumption to be reliable.

An alternative approach is to obtain confidence bounds through bootstrapping. In Stewart (2013), to address the small sample sizes often associated with QFASA, bootstrap CIs based on the zero-inflated beta distribution are presented. Given a sample of QFASA diet estimates, the function *conf*.*meth* in the *QFASA* R package returns simultaneous CIs for the true contribution of each prey type in the diet (see Stewart et al. (2025) for further details). A downside to these intervals is that they aim to estimate a common diet for a group of predators and do not therefore capture potential individual differences in diets.

Given initial MUFASA diet estimates 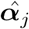 from *n* predators, we propose the following algorithm to derive bootstrap marginal confidence intervals (i.e., a CI for each prey type) for the diet of each individual in the sample:

1. Using empirical estimates of the mean prey FA signature and pooled variance-covariance matrix, and ML diagonal variance-covariance matrix of error, 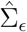 (obtained from MU-FASA), randomly sample from the parametric distributions according to Equation 4. Take the linear combination of the simulated prey FAs and MUFASA diet estimates from original predators, then perturb the resulting vector by the simulated error, as shown in Equation 3.
2. Estimate the diet proportions, 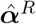, for the *n* simulated FA signatures from step 1.
3. Repeat steps 1 and 2 *R* times.
4. For each of the *n* predators, *R* parametric bootstrap replicates are available. Obtain the 2.5% and 97.5% quantiles of these replicates, one at a time, for each diet proportion (each prey) to obtain the *n* marginal CI bounds.

### 2.3 Simulation Study

We conducted simulation studies to assess the behaviour of MUFASA. A reallife prey database was required to create realistic FA signatures of pseudo-predators. The prey database we used is described below.

#### 2.3.1 Prey Database Sampling

The prey FA database used for our simulation study is a subset of the larger prey set analyzed by Budge et al. (2002). For details of sample collection and processing see Budge et al. (2002). From the larger prey set, FA signatures from 21 prey species were selected for use in simulations (Table 1).

**Table 1.** Species, sample sizes, and average lipid content (% wet weight) included in the prey database used in simulations. Asterisk (*) identifies the invertebrate species.

| Common Name | Scientific Name | $n_s$ | % lipid |
| --- | --- | --- | --- |
| American plaice | *Hippoglossoides platessoides* | 134 | 2.33 |
| Atlantic butterfish | *Peprilus triacanthus* | 75 | 10.83 |
| Atlantic cod | *Gadus morhua* | 109 | 2.46 |
| Atlantic herring | *Clupea harengus* | 229 | 6.44 |
| Atlantic mackerel | *Scomber scombrus* | 32 | 5.14 |
| Capelin | *Mallotus villosus* | 162 | 5.21 |
| Longhorn sculpin | *Clupea harengus* | 45 | 2.06 |
| Northern sandlance | *Ammodytes dubius* | 148 | 5.26 |
| *Northern shortfin squid | *Illex illecebrosus* | 35 | 3.01 |
| Pollock | *Pollachius virens* | 53 | 2.44 |
| Redfish | *Sebastes sp.* | 54 | 7.10 |
| Sea raven | *Hemitripterus americanus* | 71 | 1.97 |
| Silver hake | *Merluccius bilinearis* | 58 | 1.60 |
| Smooth skate | *Malacoraja senta* | 33 | 2.53 |
| Snake blenny | *Lumpenus lumpretaeformis* | 18 | 2.43 |
| Thorny skate | *Amblyraja radiata* | 83 | 2.59 |
| White hake | *Urophycis tenuis* | 80 | 1.29 |
| Winter flounder | *Pseudopleuronectes americanus* | 50 | 1.95 |
| Winter skate | *Leucoraja ocellata* | 40 | 1.47 |
| Witch flounder | *Glyptocephalus cynoglossus* | 24 | 1.91 |
| Yellowtail flounder | *Limanda ferruginea* | 156 | 2.21 |

FAs are described throughout by the shorthand nomenclature of A:Bn-X where A represents the carbon chain length, B the number of double bonds and X the location of the double bond nearest the terminal methyl group. For each prey, 67 FA proportions were measured. The dietary FA subset of these (as defined in Iverson et al. (2004)), excluding 16:3n-1 and 22:2n-6, was used for our simulations (Table S1 in the Supplementary Material). These 2 FAs were not consistently identified across all samples in the database and were, therefore, removed from all analyses. The remaining 29 FAs include only those FAs that are not biosynthesized by the predator.

The 21 different prey species comprising the prey database are listed in Table 1, along with the sample size and average lipid, or fat content, measured as percent of fat content relative to wet weight. Additional information regarding the selection of the prey species may be found in Stewart et al. (2022).

#### 2.3.2 Pseudo-Predators

Pseudo-predators are generated FA signatures used in place of real-life predator FA signatures (Stewart (2013), Happel et al. (2016), Bromaghin (2015)) in our simulation study to assess the estimation abilities of our model. We generated pseudo-predators parametrically and non-parametrically. Pseudo-predators generated parametrically are simulated according to model assumptions. Generation in this manner allows us to assess how our method performs in a correctly specified setting and can be helpful in identifying limitations of the algorithm due to computational difficulties, for example. Non-parametric pseudo-predators create FA signatures generated in such a way that no assumptions are being made on the distribution of the data, so results based on this method will indicate how MUFASA is performing even if model assumptions are not met.

In order to generate pseudo-predators parametrically, we use the MVN distribution, which we have assumed for our real-life prey species after transformation in our model. First, using the prey database, we estimate a mean ilr transformed FA signature for each species (we could alternatively use the median, or another measure of centre), as well as a pooled variance-covariance matrix. Using these empirical estimates, we generate one ilr transformed FA signature from the MVN distribution for each prey species. These transformed FA signatures are back transformed, and a linear combination of these signatures with the true diet is performed to obtain the FA signature of our predator on the original scale, without error.

The error terms, ***ϵ***^*ilr*^, are also generated from a MVN distribution with mean **0** and variance-covariance a diagonal matrix of 0.001. Since no predator FA signatures were available for this prey database, we could not base this on real-life values. Each error term ***ϵ***^*ilr*^ is back transformed, and a perturbation (equivalent to addition on the ilr scale) is taken between the error vector and the FA signature. This perturbation yields the FA signature of the pseudo-predator.

To generate the non-parametric pseudo-predators, a bootstrap sample of prey FA signatures is first collected from each prey species. The number of prey to resample was investigated in Bromaghin (2015), where it was found that using arbitrary sample sizes yields accurate mean FA signatures, but will affect the variability. For our simulations, the total sample size of each prey is resampled with replacement and the mean (or median) FA signatures of the bootstrapped samples are calculated for each prey species. While resampling the total sample could be underestimating the true variability in the predators, we note that the error term increases the variability. The remaining steps are the same as that for parametric pseudo-predators.

### 2.4 Simulation Settings

Due to the MUFASA procedure being computationally intensive, we opted to reduce the number of prey species included in the diets for the simulation study. We note that for one data set with 25 predators, 11 prey, 29 FAs on a single compute node with a 16 core CPU (2 x Intel Xeon Gold 6238 Cascade Lake 2.10GHz), the job wall-clock time was 1 hour 56 mins 28 sec and it used 20.20GB of memory. Therefore, we looked at the FA signatures of each prey species to select three groups of four species with varying levels of similarity. In order to choose these groups, dendrograms were plotted based on Aitchison’s, chi-squared, and KL distances of the mean FA signatures for each prey species, selecting one group with similar, one group with moderate, and one group with highly different mean FA signatures. Based on these dendrograms (Figure 1), species were selected primarily using the chi-squared distance, resulting in the sets of species shown in Table 2.

**Table 2.** Prey species selected to be included in simulations based on distances between fatty acid signatures. Group 1 species are deemed to be very different in their fatty acid signatures, Group 2 species are moderately different, and Group 3 species have similar fatty acid signatures.

| Group | Species 1 | Species 2 | Species 3 | Species 4 |
| --- | --- | --- | --- | --- |
| 1 (Very Different) | Capelin | Sea Raven | White Hake | Winter Flounder |
| 2 (Different) | Capelin | Pollock | White Hake | Redfish |
| 3 (Similar) | Atl. Butterfish | Atl. Herring | Nor. Sandlance | Redfish |

**Figure 1.**
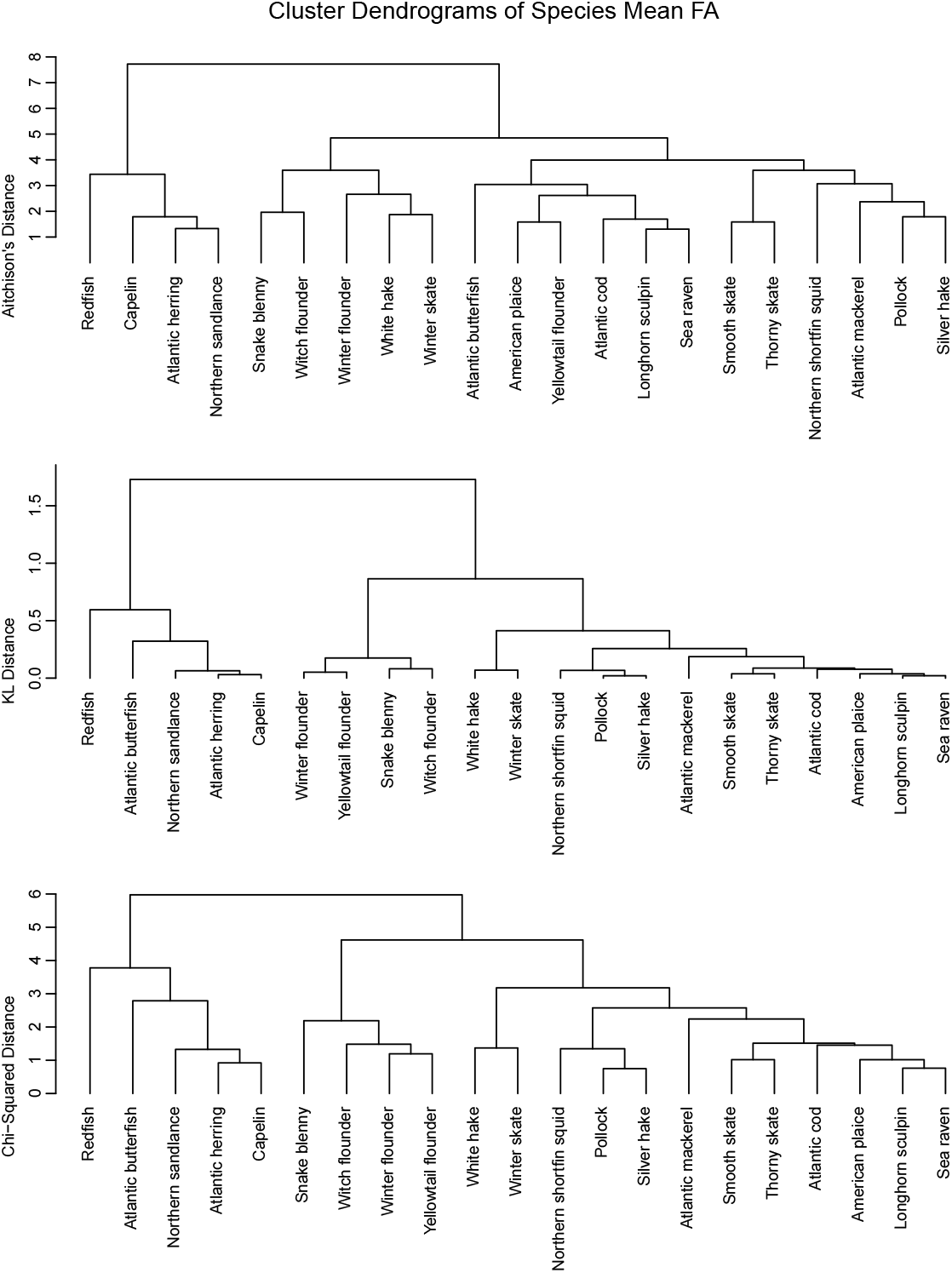
Dendrogram using Aitchison’s, KL and the chi-squared distances of the mean FA signatures for 21 prey species included in the prey data set.

For each group of species, regularly-spaced diets across the simplex were used to see how the estimates are performing throughout the whole space. We generated diets using the function *make diet grid* in the package *qfasar* (Bromaghin, 2017). Using an increment of 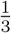, 20 diets are generated that are equally spaced by 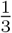 throughout the simplex. These diets ensure that the method is working not only throughout the interior of the simplex, but also on the edges. We argue that if MUFASA behaves as expected in the variety of cases that are covered by both the simulations and real-life study discussed in Section 2.5, it has the potential to work well in practice, particularly when the number of prey species is small.

Each of these diet and species combinations is run with a sample size *n* = 100 where pseudo-predators are generated randomly via both methods described, using the dietary subset of FAs. QFASA estimates (from *p*.*QFASA* in Stewart et al. (2024)) are used as starting values for the diet parameters, which were obtained using calibration coefficients set to one, KL distance, the dietary FA subset and no fat content adjustment. For *Z*, the random effects in Equation 3, a matrix of the mean ilr transformed FA signatures for each prey species is used for the starting values.

### 2.5 Real-Life Analysis

The empirical dataset used to validate MUFASA is from a captive feeding study conducted at the Vancouver Aquarium by Chad Nordstrom et al. as described in Nordstrom et al. (2008). The study, conducted between August 28 and October 9, 2003, used 21 harbour seals (*Phoca vitulina richardsi*) that were recovered from the coastline of British Columbia, Canada by the Vancouver Aquarium’s Marine Mammal Rescue Centre staff, or were brought to the rescue facility by members of the public. All seals brought to the facility were unweaned, and estimated to be less than fifteen days of age and there were no data on their feeding history. Following arrival at the facility, seals were housed in individual tubs and were tube-fed a homogenous mixture of pure salmon oil (commercial blend), ground Pacific herring (*Clupea pallasii*) and water at a ratio of 3:6:8 by weight for 5-21 days. Seals were then transitioned from the homogenate onto whole herring over a period of 5-6 days after which they received solely herring until the start of the experimental period (4-30 days). Seals were transferred to larger shared pools as they increased in size, during which time every effort was made to feed seals individually.

At the start of the experimental period (Day 0), the seals were placed in one of three diet treatments (7 seals/treatment); only Pacific herring for 42 days, only surfsmelt (*Hypomesus pretiosus*) for 42 days, or surfsmelt for 21 days followed by herring for 21 days. The seals were weighed to the nearest 0.1 kg and a full depth blubber biopsy was obtained at Day 0, Day 21 and Day 42 as described in Nordstrom et al. (2008). Not all seals were available for biopsy at Day 42. Four seals in the herring diet group were released without biopsy 34 days after the start of the experimental diets (after completing the rescue program) resulting in a reduced sample size for this group.

This diet treatment which switched from surf smelt to herring after 21 days had recognized shortcomings. Most notably, Nordstrom et al. (2008) had trouble with the prey intake by individuals in this treatment group and, as a result, individuals in this group gained considerably less mass than in the other two treatment groups over the 42 day treatment period. Consequently, the expected/predicted contributions of the prey fed to the diet were not well estimated by QFASA in Nordstrom et al. (2008) (high variability in estimates, significant overestimation of herring and underestimation of smelt intake). Given the logistical challenges with this feeding trial and the associated challenges with estimating expected contributions to the blubber signature in animals with occasional periods of weight loss and overall poor weight gain in the absence of body composition data, we did not consider it a good representation of animals eating a known mixed diet and it was removed from our consideration.

Daily food intake (to the nearest 0.01 kg) was recorded for all individuals throughout the study. Individual whole prey were randomly subsampled throughout the study period and stored in airtight plastic bags frozen at -20^*◦*^C until analysis (Table 3). The surfsmelt fed during the study were comprised of two size classes: small (9.7-11.4 cm fork length, 9.5-15.5 g wet weight) and large (13.6-17.1 cm fork length, 24.5-53.2 g wet weight). Subsamples were taken from both sizes of smelt for subsequent analysis (Table 3). In addition to the three prey items fed to the seals (salmon oil, herring and smelt), whole individuals were obtained for eight other species of fish and invertebrates and stored in airtight plastic bags frozen at -20^*◦*^C until analysis. Prey specimen and seal blubber biopsies (Nordstrom et al., 2008) were analysed using techniques described in Budge et al. (2006) and Iverson et al. (1997).

**Table 3.** Sample sizes and average lipid (%) of the 11 species (or 12 prey groups since Surfsmelt has been divided by size into two groups) included in the prey database for harbour seals.

| Common Name | Scientific Name | $n_s$ | Average Lipid (%) |
| --- | --- | --- | --- |
| Capelin | *Mallotus villosus* | 54 | 3.28 |
| Coho | *Oncorhynchus kisutch* | 38 | 3.79 |
| Eulachon | *Thaleichthys pacificus* | 30 | 8.80 |
| Herring | *Clupea harengus* | 23 | 11.23 |
| Mackerel | *Scomber scombrus* | 24 | 5.87 |
| Pilchard | *Sardina pilchardus* | 18 | 20.33 |
| Pollock | *Pollachius virens* | 17 | 6.88 |
| Salmonoil | *Salmo salar* | 5 | 100.00 |
| Sandlance | *Ammodytes dubius* | 15 | 5.73 |
| Squid | *Teuthida* | 43 | 2.21 |
| Surfsmelt lg | *Hypomesus pretiosus* | 30 | 2.85 |
| Surfsmelt sm | *Hypomesus pretiosus* | 10 | 2.24 |

In order to assess model performance, diet estimates computed from MUFASA (using the dietary FA list excluding FA 16:4n-3) were compared to the true known diets from the capture study described above. In Nordstrom et al. (2008), the true diets were calculated using a cumulative recorded diet intake for each individual over the assessment period between 35 and 75 days prior to each of the biopsy periods. This method ignores the presence of existing fat stores that the animals may have had when they arrived at the facility and assumes that there is complete or almost complete turnover of FAs. To account for the contribution of the blubber present at the time of admission to the overall blubber FA signature and to obtain estimates for Days 0, 21 and 42, we calculated the estimated contribution of each prey species to the blubber FA signature of each individual. For each study animal, the cumulative mass intake of the prey fed was used to estimate the expected contribution of each prey species to the blubber fatty acid signature (as a proportion of diet) at Days 0, 21, and 42 of the experimental period. The body fat content of the study animals at the time of admission to the rescue facility was not known. Therefore, to account for the contribution of the blubber present at the time of admission to the overall blubber FA signature at each experimental time point, a baseline body fat content at admission was estimated, and then it was assumed that the FAs subsequently consumed were deposited with existing fat, and used as a single pool (see Iverson et al. (2004)). Because the study animals were growing pups, it was also assumed that a fraction of the fatty acids consumed were immediately oxidized and not deposited. For all individuals, the amount of blubber present at the time of admission was estimated using the value for newborn harbour seal pups from Bowen et al. (1992) of 11.3% body fat. This value may be an overestimate for pups which may have lost significant body condition following the separation from their mother but it provided a reasonably conservative estimate in the absence of information on prior feeding history (i.e. duration of suckling before separation), the subsequent duration of separation, or body condition at the time of arrival to the facility (condition score or condition index; Lander et al. (2003)). Storage efficiency (the proportion of energy intake subsequently deposited) has not been examined for harbour seal pups, therefore, the value of 70% obtained for nursing pups in the closely related grey seal (*Halichoerus grypus*; Mellish et al. (1999), Lang et al. (2011)) was used.

To obtain species-specific calibration coefficients, full depth blubber biopsies were taken as described in Nordstrom et al. (2008) from 4 captive sub-adult harbour seals (3 males, 1 female) housed at the Vancouver Aquarium. These individuals had exclusively eaten herring from a single lot (the same lot used in the experimental diet treatments above) for *>*1 yr prior to the biopsies. The calibration coefficients used in the modelling were calculated as the 10% trimmed mean using the FA signatures from these 4 seals and 23 herring subsampled over the feeding period following the method of Iverson et al. (2004).

## 3 Results

### 3.1 Simulations

To measure how well MUFASA performed relative to QFASA, we computed the chi-square distance proposed in Stewart (2017). The chi-square distance is beneficial as it does not require modification of the zeros in the estimates or true diets. For each of the 100 pseudo-predators in all 20 diets explored, the chi-squared distance was calculated between the estimates (both QFASA and MUFASA) and the true diet. These distances for species group 1 based on pseudo-predators generated parameterically, are summarized in Figure 2, where we see that nearly all of the QFASA estimates appear to be much further from the true diet than MUFASA.

**Figure 2.**
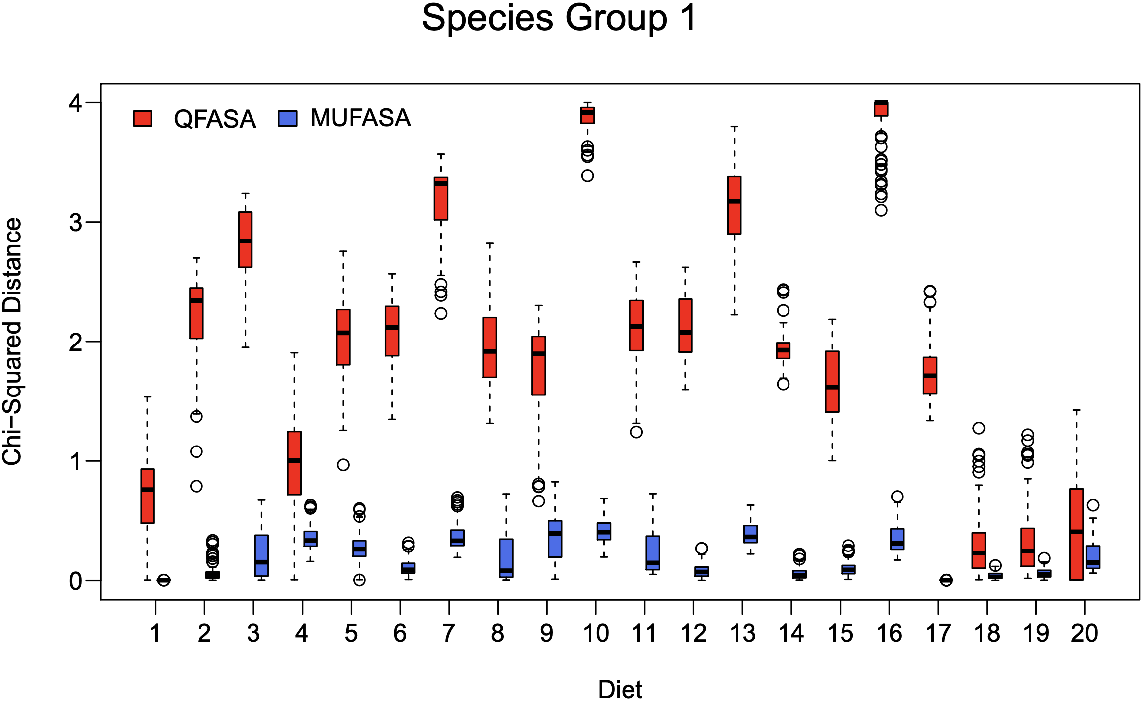
Boxplots of the 100 chi-squared distances for each of the 20 diet simulations, using parametrically generated pseudo predators and species in Group 1. That is, species that are deemed to be very different in their fatty acid signatures.

To examine the behaviour of the diet estimates more closely on an individual basis, we selected two diets informed by Figure 2: one that appeared to be more difficult to estimate (as MUFASA and MLE estimates differed between the two methods) and one that was less problematic (produced similar estimates). Since these diets corresponded to diets 16 and 18 respectively in our simulations, and they are denoted in this way in the boxplots in Figures 4 and 5 displaying the corresponding diet estimates, where the purple lines represent the true proportions in each of the diets.

**Figure 3.**
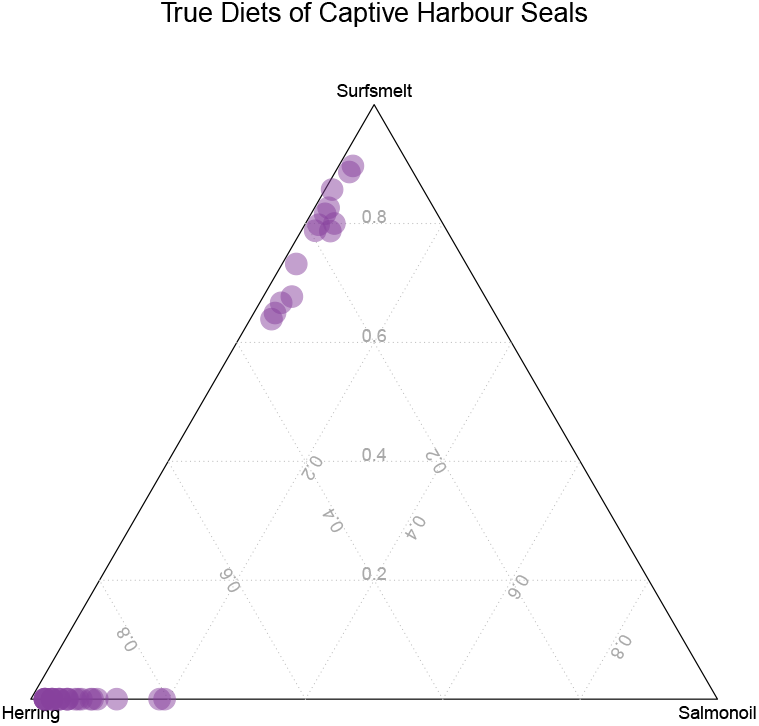
Ternary diagram of the true diets of 38 biopsies of harbour seals during a captive study at Vancouver Aquarium. Points along the edges indicate diets where the prey species at the opposing vertex is not consumed, with the diet composed only of the other two prey species.

**Figure 4.**
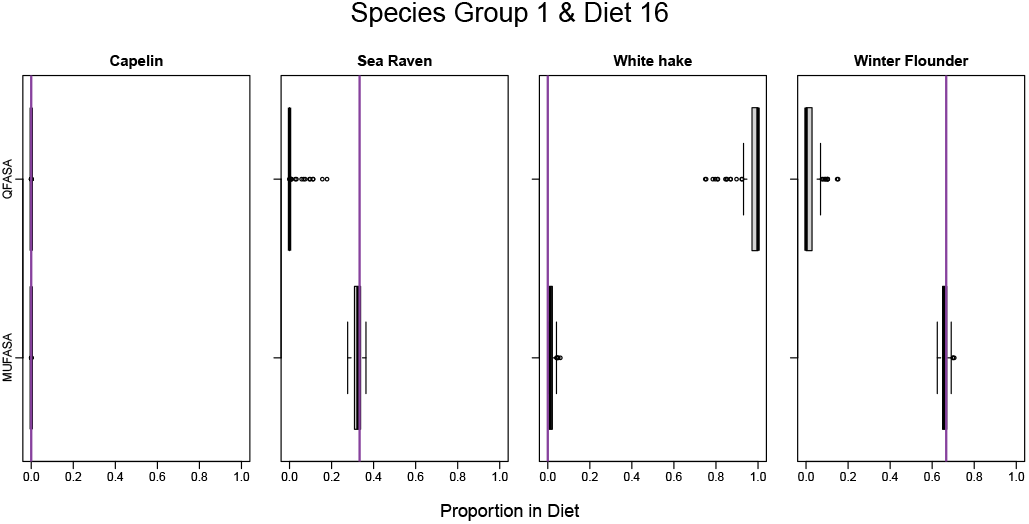
Boxplots of estimated diet proportions of 100 parametric pseudo-predators using species in Group 1 (that is, species that are deemed to be very different in their fatty acid signatures) and diet 16 for both QFASA and MUFASA, where the true diet is shown in purple.

**Figure 5.**
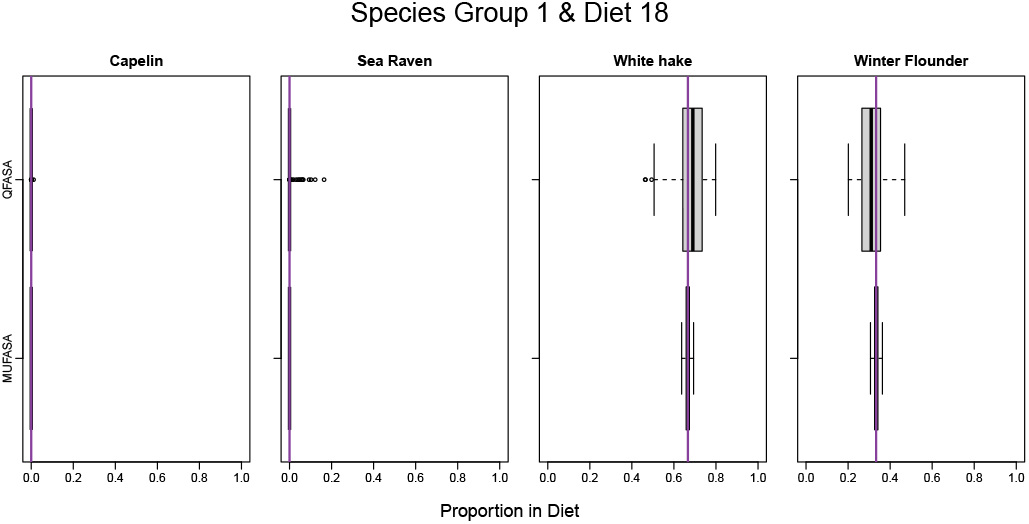
Boxplots of estimated diet proportions of 100 parametric pseudo-predators using species in Group 1 (that is, species that are deemed to be very different in their fatty acid signatures) and diet 18 for both QFASA and MUFASA, where the true diet is shown in purple.

For diet 16, shown in Figure 4, the true diet (0, 0.333, 0, 0.667) is located inward of the edges of the simplex. QFASA overestimated white hake, and underestimated winter flounder and sea raven. Further analysis using the ‘additional measures’ output from *p*.*QFASA* (in the *QFASA* R package), show that FAs 20:5n-3, 16:3n-4, and 16:3n-6 contribute the most on average to the distance used to estimate the diets. Together, they account for over 45% of the distance between the predator FA signature and the linear combination. The FA 20:5n-3 is present in the largest proportions in the predator signatures so it makes sense that 20:5n-3 contributes a lot to the distance. However, 16:3n-4 and 16:3n-6 have relatively small proportions in the predator signatures, as they are just slightly larger than the first quartile. Those same FAs did not appear to affect MUFASA as much as with QFASA. We hypothesize that MUFASA does not inflate the contribution of these FAs, which results in more accurate estimates in this case.

The results for species groups 2 and 3 were similarly explored. As with species group 1, in nearly all cases, MUFASA yielded similar estimates, if not more accurate and precise, than the original QFASA method. As results were quite similar, it was decided that it would be redundant to show all plots, but readers may see Steeves (2020) for additional results.

Similarly, simulations using pseudo-predators generated non-parametrically were conducted by way of resampling prey FA signatures from the prey database and taking a linear combination using the same settings as in the parametric case. As findings were similar for the 3 groups (see Steeves (2020)), only species group 1 is reported. As expected, the MUFASA estimates from these simulations were not as accurate as those from parametric simulations (see Figure 6).

**Figure 6.**
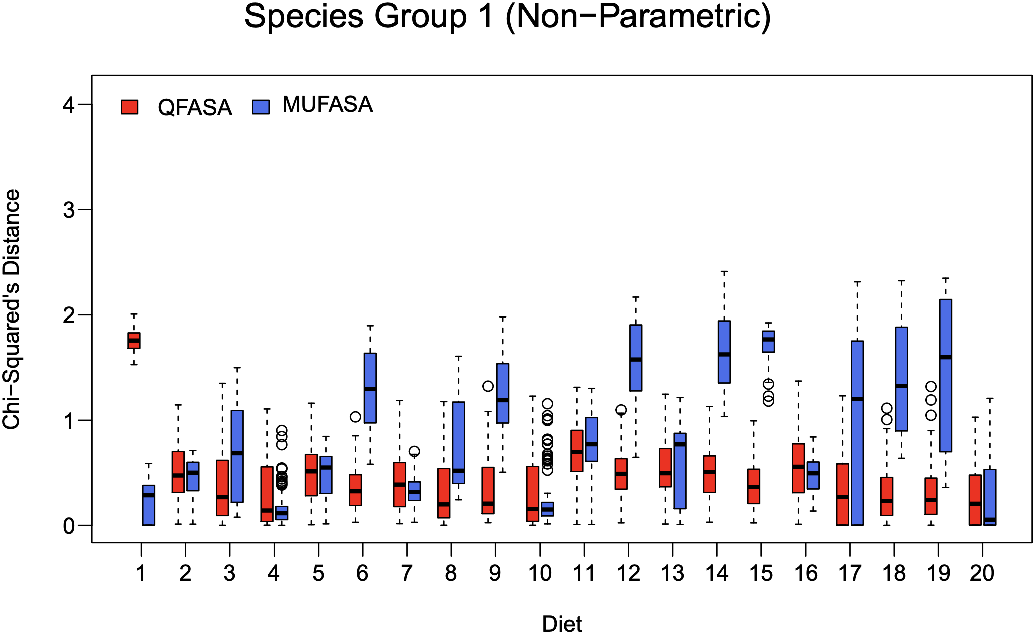
Boxplots of the 100 chi-squared distances for each of the 20 diet simulations, using non-parametrically generated pseudo predators and species in Group 1. That is, species that are deemed to be very different in their fatty acid signatures.

In the nonparametric setting, MUFASA yielded estimates further from the true diet than QFASA in some cases, and often with less precision. For example, diet 15 has large differences between the true diet and estimated diet for MUFASA. The boxplot depicting diet 15 estimates is displayed in Figure 7). For all species, QFASA is estimating the diet relatively accurately with a range up to 10% below and 10% above the true values. With MUFASA, Capelin is only slightly overestimated, sea raven is overestimated between 0 and 15%, white hake is underestimated by approximately 30%, and winter flounder is overestimated by between 0 and 10%. Despite being one of the worst estimations of the non-parametric simulations using MUFASA, the differences are not nearly as large as in the parametric simulations for QFASA.

**Figure 7.**
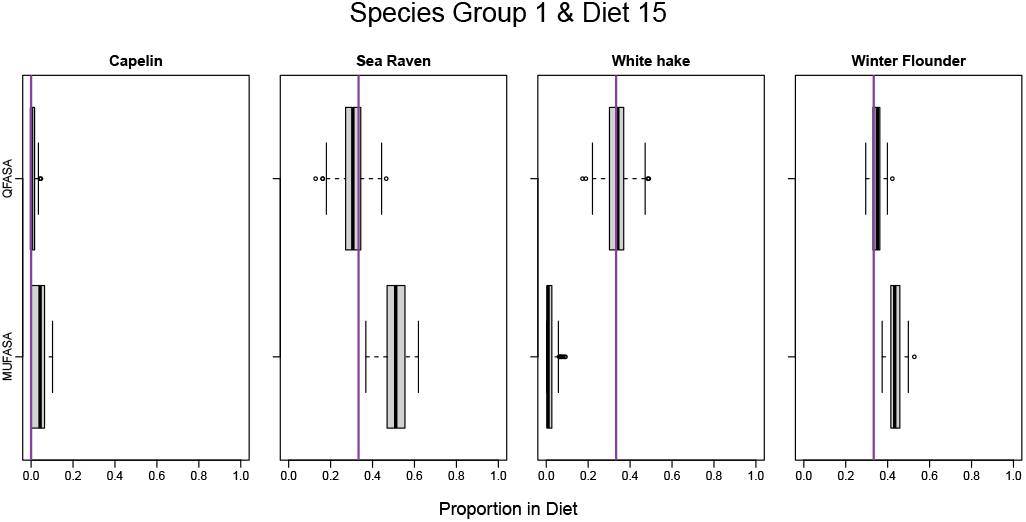
Boxplots of estimated diet proportions of 100 non-parametric pseudo-predators using species Group 1 and diet 15 for both QFASA and MUFASA, where the true diet is shown in purple.

Similarly, we can see from Figure 6 that diet 1 has relatively small differences between the true proportions and the estimated proportions of MUFASA. Diet 1 estimates are displayed in a boxplot in Figure 8. Here, we see that MUFASA is estimating all species accurately and with slightly smaller variability than QFASA. QFASA seems to be underestimating Capelin while overestimating Sea Raven.

**Figure 8.**
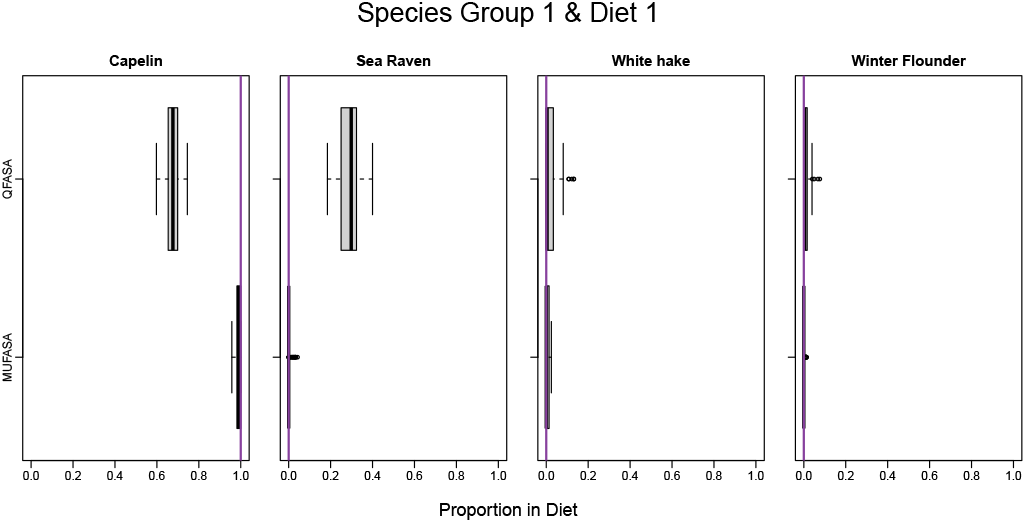
Boxplots of estimated diet proportions of 100 non-parametric pseudo-predators using species Group 1 and diet 1 for both QFASA and MUFASA, where the true diet is shown in purple.

Results of a limited simulation study carried out to investigate the coverage probabilities and interval lengths associated with the proposed the bootstrap CI algorithm may be found in the Supplementary Material. Due to the computational demands of the MUFASA algorithm itself, combined with the bootstrapping procedure, it was necessary to constrain the scale of our simulation study. Preliminary results, however, suggest that the proposed CI method generally works well except for when the true diet is 0 or 1. This is because when the true diet proportion is on these bounds, it is impossible to obtain a confidence bound below 0, or above 1, using the percentile method. However, when sufficiently rounded, the limits do include 0 or 1 as desired.

### 3.2 Captive Harbour Seals

The captive harbour seal study conducted at Vancouver Aquarium (Nordstrom et al., 2008) provides an opportunity to compare the true parameters of interest with the estimated values. Thirty-eight true diets were obtained as described in Section 2.5 and are displayed in a ternary diagram shown in Figure 3. Note that in Figure 3 we only include the prey items that were fed to the seals, namely herring, salmon oil and surfsmelt. The ternary diagram shows that the diets mostly contain herring or surfsmelt, as most of the dots are very close to, if not on the edges. Further, the 38 biopsies (and subsequent diet estimates) are not independent as they are derived from 14 distinct seals so, for this reason, we did not compute CIs using this data set.

In this analysis, calibration coefficients and fat content described in Section 2.5 were implemented using standard techniques. Since the diet of each individual seal was different, instead of plotting the diet estimates, the bias of each individual (true diet minus estimated diet), is plotted in a box plot shown in Figure 9. Note that the scale is not the same for all species in this figure. From Figure 9, we can see that for most species that have true diet proportion 0, namely, capelin, coho, eulachon, mackerel and pollock, both QFASA and MUFASA are performing quite well, as the median bias is 0, with little to no variation. For other species with true proportion zero, namely pilchard, sandlance, and squid, both methods have difficulty estimating zero. Both QFASA and MUFASA underestimate all three of these, pilchard by about 10%, sandlance by about 5% and squid by around 5%. In general, MUFASA and QFASA produced similar estimates with relatively small biases on average.

**Figure 9.**
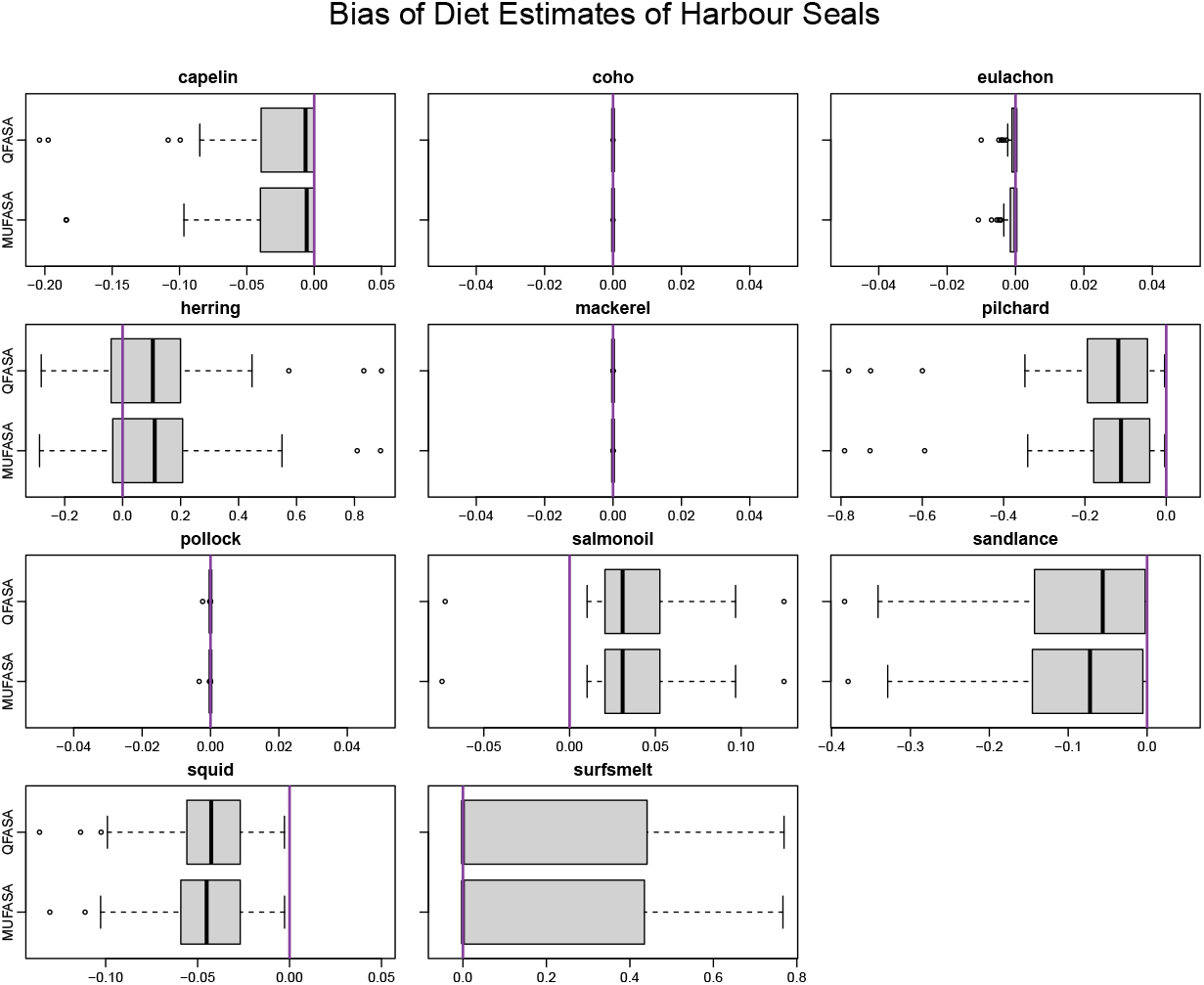
Bias (estimated diet - true diet) for 38 harbour seals, estimated using both MU-FASA and QFASA.

To better understand the causes of the observed biases, we compared the FA signatures of the species using the dendrogram shown in Figure 10. Prey species with similar FA signatures are difficult to differentiate from each other, and thus could explain over or under estimation. For example, pilchard and herring have very similar FA signatures, and we can see in Figure 9 that herring is overestimated by approximately 10% and pilchard is underestimated by approximately 10%. Therefore, the similarities in FA signatures may be creating a slight bias in estimation for these two species. Similar situations may arise between surfsmelt and sandlance. There is a large amount of variability in the bias of surfsmelt estimates. This may be due to confounding with sandlance, particularly for smaller individuals. This is consistent with findings in Nordstrom et al. (2008), where surfsmelt was often overestimated.

**Figure 10.**
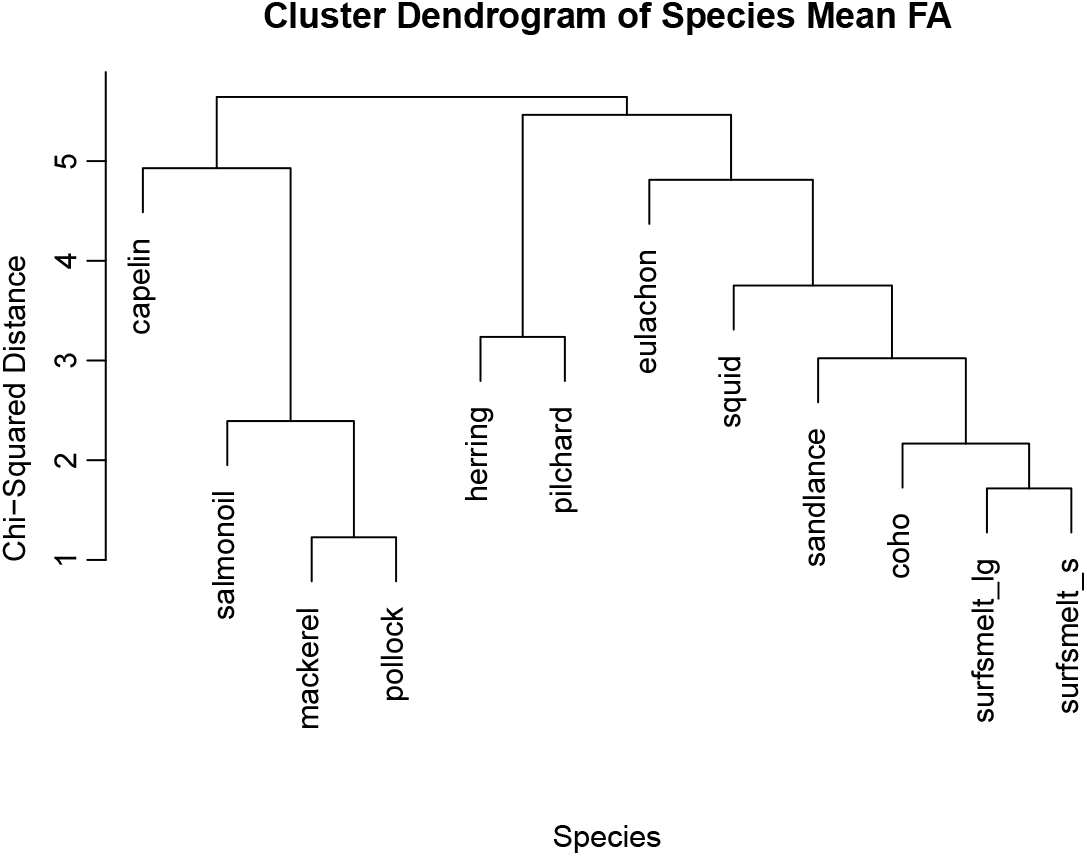
Dendrogram of 11 prey species included in the Nordstrom prey database, using the mean FA signatures, and the chi-squared distances.

## 4 Discussion

Developing a ML estimation method for diet proportions of predators based on QFASA required careful consideration of what is happening both biologically and statistically. The weighted mixture of diet proportions and prey FA signatures could be considered in two spaces: the simplex, where we would take a mixture of FA signatures on the original scale, or real Euclidean space, where we would take a mixture of transformed FA signatures. After considering the biological implications of both techniques, we decided that taking the mixture on the original scale was the most reasonable, as the FAs would be absorbed into the predator tissues without scaling. As well, since the actual prey consumed by the predator would never be included in the sample, these unobserved prey were represented by a random effect. Our model assumes that after transformation, the predator and prey data are multivariate normally distributed. Our results suggest that MUFASA is relatively robust; in practice the MVN assumption should be investigated. In Steeves (2020), removal of potential outliers in the FAs resulted in data that were more closely normally distributed.

Together the random effects, unknown mean vectors and variance-covariance matrices of the prey FA signatures constitute a large number of unknown parameters and has the potential to cause identifiability problems.. The mean vectors and covariance matrices for each species comprise a large percentage of the parameters. These parameters are not of primary interest in estimating diet and, provided we have reliable data for each species, we can opt to use empirical estimates. The use of empirical estimates for the mean/covariance matrices and a diagonal covariance matrix for the error also reduced the computational complexities. If the empirical estimates are of poor quality, we could potentially handle this situation by first estimating the diet proportions and fixing the means and variance-covariance matrices as we have done, then, while fixing the diet proportions obtained, estimate the mean and variance-covariance matrices. This could be repeated until there are no changes from iteration to iteration. In order to accomplish this, a higher level of computational efficiency is needed.

The simulation studies were developed and run to determine how MUFASA behaved in practice, but with some known limitations. Calibration coefficients and fat content were ignored in the simulation study and these could have an effect on the accuracy and variability of the estimates. The covariance matrix of the error terms was, to a degree, arbitrarily chosen and, if set too small, could yield FA signatures that are less variable than those sampled in practice. Using only 4 prey species in the diet for a predator, specifically grey and harbour seals, is not realistic and ideally a simulation study using more prey species is desired; however, this poses a challenge for this computationally intensive model. We note that MUFASA performed well in the real-life example where calibration coefficients and fat content were included in the analysis and the prey database contained 11 species. Further exploration with additional real-life data sets are needed to assess whether our simulation assumptions are realistic in a variety of situations, and more validation studies are needed to understand the advantages and limitations of MUFASA.

MUFASA may offer an advantage over current Bayesian methods. For instance, Litmanen et al. (2020) suggests the use of QFASA over Bayesian methods with the caveat that there is no way to incorporate priors, model uncertainty, and the diet estimates are point estimates. MUFASA offers improvements in two of these areas: modelling prey uncertainty and providing a framework for inference.

## 5 Conclusion

In this article, a novel approach to quantitatively estimating diet proportions of predators using a ML approach titled MUFASA was presented. The likelihood is based on a linear combination of multivariate normally distributed random effects representing ilr transformed unobserved FA signatures of prey species, that is perturbed by a random error vector. By way of simulations, we have shown that MUFASA generally performed better than the widely accepted QFASA method when model assumptions are met. Using non-parametrically generated pseudo-data, while QFASA often performed better than MUFASA for most diets, results were comparable, implying that the new methodology has the potential to work reasonably well, even when distributional assumptions, such as MVN, are in question. We note that the diets examined were chosen from a grid and we have not explored whether better performance was associated with more realistic diets, or vice versa.

A method of obtaining individual bootstrapped confidence intervals for the diet was proposed with preliminary results suggesting that the algorithm works well when the actual diet is not 0 or 1. Diets on the edges of the simplex are less likely to be constrained in the marginal CIs. A larger study of the coverage probabilities is an area for future work, to better explore the behaviour of these CIs.

Finally, MUFASA was evaluated on a real-life study of captive harbour seals at the Vancouver Aquarium (Nordstrom et al., 2008) for which we had knowledge of the true diets. With this data, we were able to ensure that MUFASA is capable of adequately identifying and quantifying prey species in realistic FA signatures of predators, with a larger number of prey species included in the model.

As a newly developed approach, MUFASA has not yet been validated to the same extent as QFASA. However, we have shown through both simulations and a real-life application that MUFASA estimates tend to be of similar quality to those derived from QFASA in the settings explored, but with several significant advantages including a model that may better reflect the biological situation at hand with the inclusion of random effects, random errors, and modelling of the prey FA signatures. Further, the availability of a likelihood function enables the use of a broad range of existing methodologies that may have the capacity to address current well-known challenges associated with diet estimation via fatty acids.

